# Metabolic Network Analysis and Metatranscriptomics Reveals Auxotrophies and Nutrient Sources of the Cosmopolitan Freshwater Microbial Lineage aci

**DOI:** 10.1101/106856

**Authors:** Joshua J. Hamilton, Sarahi L. Garcia, Brittany S. Brown, Ben O. Oyserman, Francisco Moya-Flores, Stefan Bertilsson, Rex R. Malmstrom, Katrina T. Forest, Katherine D. McMahon

## Abstract

An explosion in the number of available genome sequences obtained through metagenomics and single-cell genomics has enabled a new view of the diversity of microbial life, yet we know surprisingly little about how microbes interact with each other or their environment. In fact, the majority of microbial species remain uncultivated, while our perception of their ecological niches is based on reconstruction of their metabolic potential. In this work, we demonstrate how the “seed set framework”, which computes the set of compounds that an organism must acquire from its environment (1), enables computational analysis of metabolic reconstructions, while providing new insights into a microbe’s metabolic capabilities, such as nutrient use and auxotrophies. We apply this framework to members of the ubiquitous freshwater Actinobacterial lineage acI, confirming and extending previous experimental and genomic observations implying that acI bacteria are heterotrophs reliant on peptides and saccharides. We also present the first metatranscriptomic study of the acI lineage, revealing high expression of transport proteins and the light-harvesting protein actinorhodopsin. Putative transport proteins complement predictions of nutrients and essential metabolites while providing additional support to the hypothesis that members of the acI are photoheterotrophs.

**Importance:** The metabolic activity of uncultivated microorganisms contributes to numerous ecosystem processes, ranging from nutrient cycling in the environment to influencing human health and disease. Advances in sequencing technology have enabled the assembly of genomes for these microorganisms, but our ability to generate reference genomes far outstrips our ability to analyze them. Common approaches to analyzing microbial metabolism require reconstructing the entirety of an organism’s metabolic pathways, or performing targeted searches for genes involved in a specific process. This paper presents a third approach, in which draft metabolic reconstructions are used to identify compounds through which an organism may interact with its environment. These compounds can then guide more intensive metabolic reconstruction efforts, and also provide new hypotheses about the specific contributions microbes make to ecosystem-scale metabolic processes.

## Introduction

Natural microbial communities have central roles in the biosphere, ranging from mediators of nutrient cycling to agents of human health and disease (2, 3). However, the majority of microbial species remain uncultivated, a feature that poses a significant challenge to our understanding of their physiology and metabolism. Recent advances in sequencing technology and bioinformatics have enabled assembly and analysis of reference genomes for a wide range of hitherto uncultured community members from diverse environments (4) that can be used to reconstruct an organism’s metabolism.

Common approaches to metabolic reconstruction involve the comprehensive reconstruction of an organism’s metabolic pathways (5), or a targeted search for genes involved in processes of interest (6). These reconstructions can then be analyzed manually or using computational approaches such as flux-balance analysis (FBA) (7). However, FBA-based approaches require a comprehensive understanding of an organism’s growth requirements and biomass composition, information which is often unavailable for uncultivated microorganisms. An alternative approach is to compute an organism’s *seed set*, the set of compounds that the organism cannot synthesize on its own and must exogenously acquire from its environment (e.g., its growth requirements) (1). These compounds may represent both *auxotrophies*, essential metabolites for which biosynthetic routes are missing, and *nutrients*, compounds for which degradation but not synthesis routes are present in the genome. The *seed set framework* offers potential advantages over other reconstruction-based approaches, as identification of seed compounds facilitates a focused analysis by identifying those compounds through which an organism interacts with its environment.

In the present study, we present a computational pipeline to predict seed compounds using metabolic network reconstructions generated from KBase (8). We apply this pipeline to a collection of 36 metagenome-assembled genomes (MAGs) and single-cell genomes (SAGs) from the abundant and ubiquitous freshwater Actinobacterial lineage acI, which is thought to have a central role in nutrient cycling in diverse freshwater systems (9–18). The seed compounds predicted by our analysis are in agreement with previous experimental and genomic observations (19–27), confirming the ability of our method to predict an organism’s auxotrophies and nutrient sources.

In particular, we find that members of the acI lineage are auxotrophic for essential vitamins and amino acids, and may consume as nutrients a wide array of N-containing compounds (including ammonium, branched-chain amino acids, polyamines, and di- and oligo-peptides) as well as mono-, poly-, and oligo-saccharides. To complement these predictions, and to understand which pathways dominate active metabolism of acI in its natural environment, we conducted an *in situ* metatranscriptomic analysis of gene expression in the acI lineage. This analysis revealed that the acI express a diverse array of transporters for auxotrophies, nutrients, and other compounds that may contribute to their observed dominance and widespread distribution in a variety of aquatic systems.

## Results

### Phylogenetic Affiliation of acI Genomes

We identified 17 SAGs and 19 MAGs from members of the acI lineage (Supplementary Table S1) in a larger set of reference genomes derived from our long-term study sites. A phylogenetic tree of these genomes built using a concatenated alignment of single-copy marker genes is shown in Figure 1. Previous phylogenetic analyses using 16S rRNA gene sequences showed that the acI lineage can be grouped into three distinct monophyletic clades (acI-A, acI-B, and acI-C) and thirteen so-called “tribes” (28). In this study, the phylogenetic tree also identified three monophyletic branches, enabling MAGs to be classified to clade and tribe level based on the taxonomy of SAGs within each branch (as determined by 16S rRNA gene sequences, either PCR amplified or assembled from the single cell). Note that three MAGs formed a monophyletic group separate from clades acI-A and acI-B; we assume these genomes belong to clade acI-C as no other acI clades have been identified to date.

**Figure 1.**
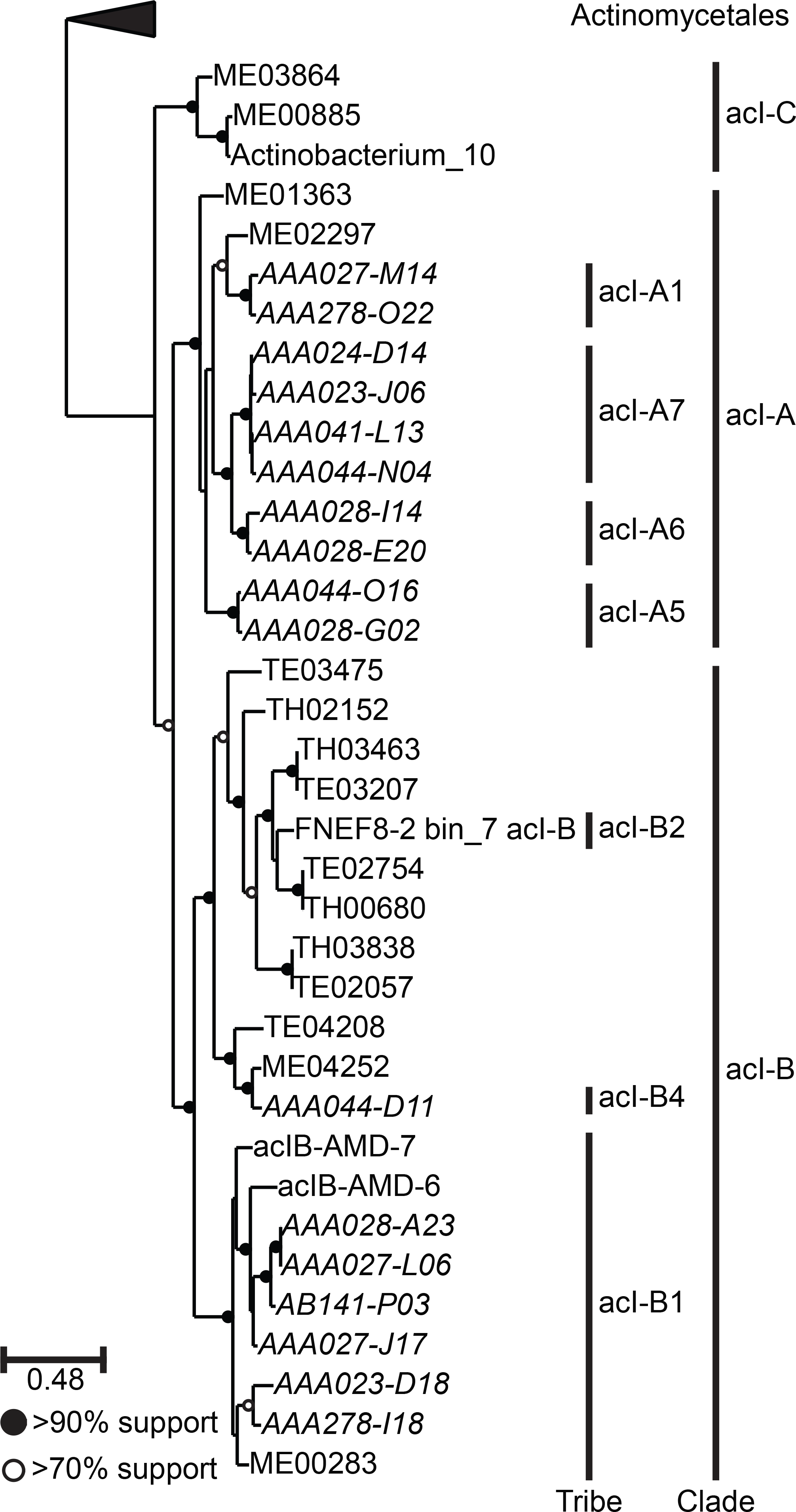
Phylogenetic placement of the genomes used in this study within the acI lineage. The tree was built using RAxML (40) from a concatenated alignment of protein sequences from 37 single-copy marker genes (39). The order Actinomycetales forms the outgroup. Vertical black bars indicate groups of genomes belonging to defined tribes/clades within the acI lineage, as determined using 16S rRNA gene sequences (for SAGs and bin FNEF8-2 bin_7 acI-B only) and a defined taxonomy (28). SAGs are indicated with italic text. Supplementary Figure S1 shows the position of the acI lineage relative to other orders within the class Actinobacteria.

### Estimated Completeness of Tribe- and Clade-Level Composite Genomes

We constructed composite genomes from multiple SAGs and/or MAGs, to partially alleviate limitations presented by incomplete genomes. To do this, we first estimated the completeness of tribe- and clade-level composite genomes using singlecopy marker genes. This allowed us to determine the finest level of taxonomic resolution at which we could confidently compute seed compounds, using genome completeness as a proxy for metabolic reaction network completeness (Figure S2). We deemed genomes to be nearly complete if they contained 95% of the lineage-specific marker genes. With the exception of tribe acI-B1, tribe-level composite genomes are estimated to be incomplete (Figure S2A). At the clade level, clades acI-A and acI-B are estimated to be nearly complete, while the acI-C composite genome remains incomplete, as it only contains 75% of the 204 marker genes (Figure S2B). As a result, seed compounds were calculated for composite clade-level genomes, with the understanding that some true seed compounds for the acI-C clade will not be predicted.

### Computation and Evaluation of Potential Seed Compounds

Metabolic network reconstructions for each genome were built using KBase. Composite metabolic network graphs were then constructed for each tribe and clade by merging metabolic network reconstructions of individual genomes. Seed compounds for each clade were then computed from that clade’s composite metabolic network graph using a custom implementation of the seed set framework (Figure 2). A total of 125 unique seed compounds were identified across the three clades (Table S2).

**Figure 2.**
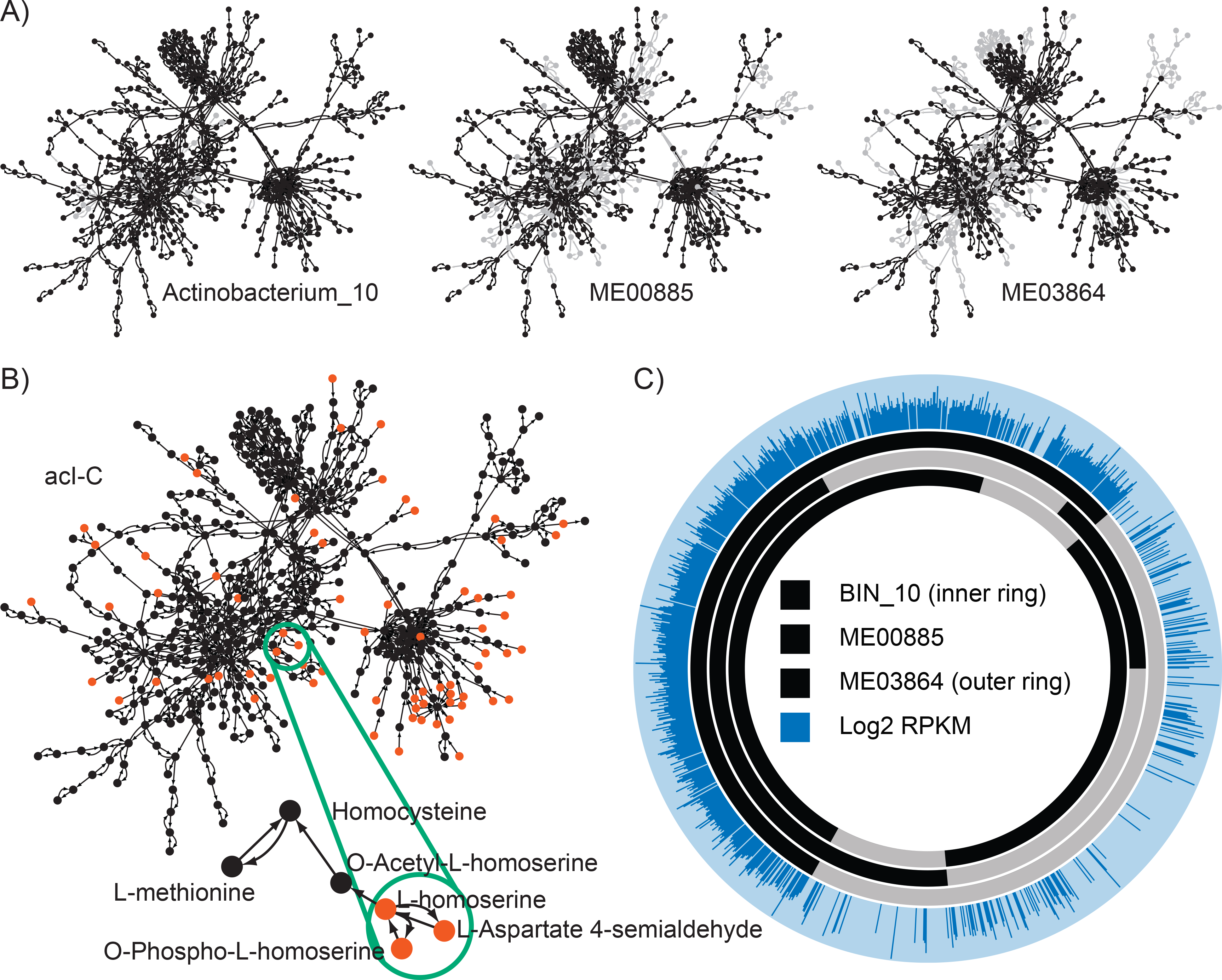
Overview of the seed set framework and metatranscriptomic mapping, using three genomes from the acI-C clade as an example. **(A)** Metabolic network graphs are created for each genome belonging to clade acI-C. In these graphs, metabolites are represented as nodes (circles) and reactions by arcs (arrows). Grey nodes and edges indicate components of the composite graph missing from that genome graph. Additional information on this step of the workflow is available in Figure S2. **(B)** A composite network graph is created for each clade by joining graphs for all genomes from that clade, and seed compounds (red) are computed for the composite graph. Additional information on this step of the workflow is available in Figures S3, S4, and S5. **(Inset)** Three seed compounds which indicate an auxotrophy for L-homoserine, a methionine precursor. **(C)** Metatranscriptomic reads are mapped to each individual genome using BBMap. Orthologous gene clusters are identified using OrthoMCL (29). For each cluster, unique reads which map to any gene within that cluster are counted using HTSeq (47). The relative gene expression is computed using RPKM (48).

Because KBase is an automated annotation pipeline, the predicted set of seed compounds is likely to contain inaccuracies (e.g., due to missing or incorrect annotations). As a result, we screened the set of predicted seed compounds to identify those that represented biologically plausible auxotrophies and nutrients, and manually curated this subset to obtain a final set of auxotrophies and nutrient sources. Of 125 unique compounds, 31 (79%) were retained in the final set of proposed auxotrophies and nutrients. Tables S3 and S4 contain this final set of compounds for clades acI-A, acI-B, and acI-C, and Figure 3 shows the auxotrophies and nutrients these compounds represent.

**Figure 3.**
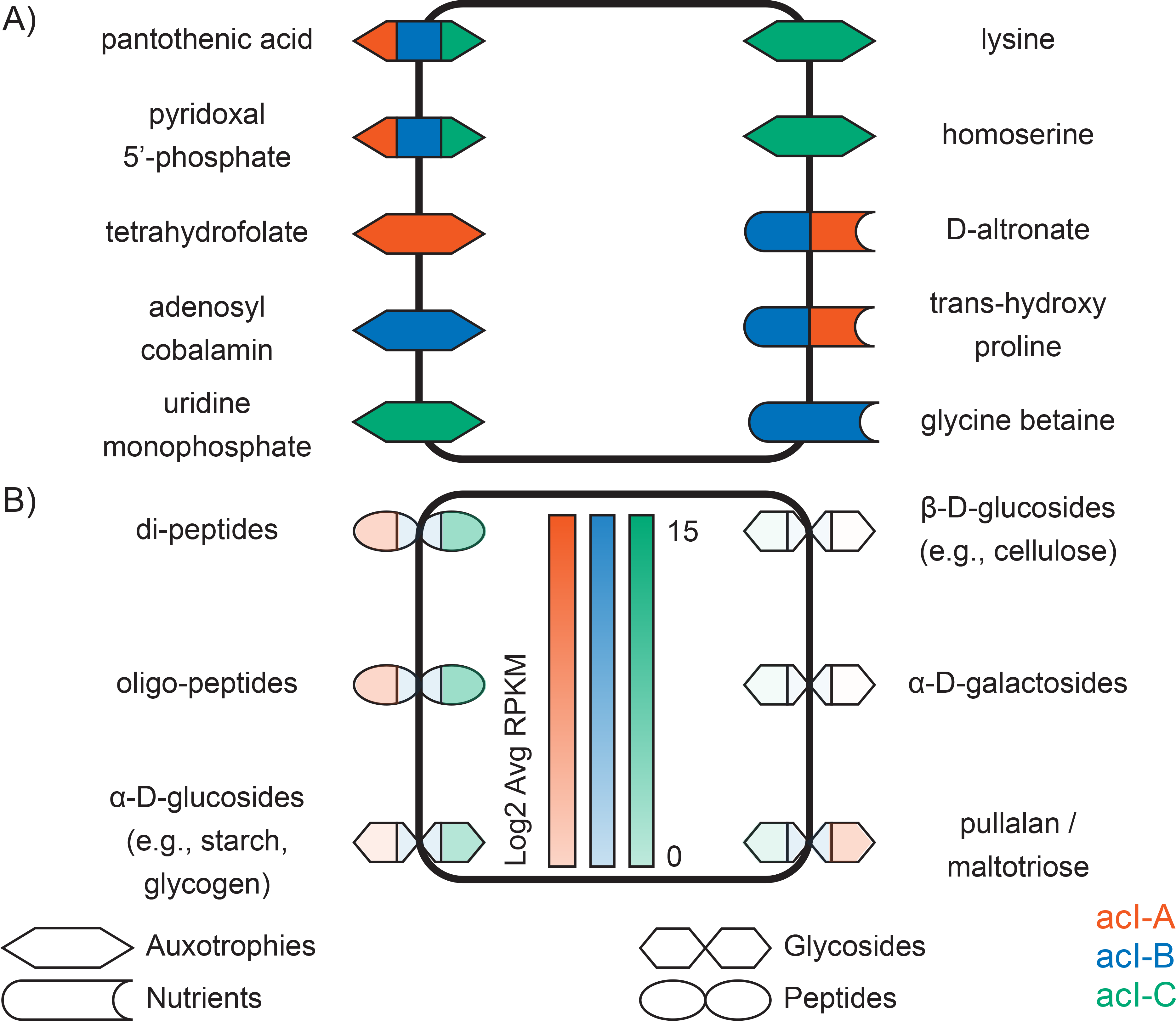
Seed compounds of members of the acI lineage. **(A)** Auxotrophies and nutrient sources, not including peptides and glycosides. **(B)** Peptides and glycosides. These compounds represent those inferred from genome annotations, rather than the seed compounds themselves. In panel (B), the intensity of the color indicates the log2 RPKM of the encoding gene cluster. For compounds acted upon by multiple gene clusters, the percentile of the most highly-expressed cluster was chosen.

### Making Sense of Seed Compounds via Protein Clustering and Metatranscriptomic Mapping

For seed compounds representing nutrient sources, genes associated with the consumption of these compounds should be expressed. To test this, we collected and sequenced four metatranscriptome samples from Lake Mendota (Dane County, WI, USA). However, because seed compounds were computed from each clade’s composite metabolic network graph, genes associated with the consumption of seed compounds may be present in multiple genomes within the clade. To facilitate the linkage of metatranscriptome measurements to seed compounds, we mapped the metatranscriptome reads to clusters of orthologous groups (COGs) within each clade. We used OrthoMCL (29) to identify COGs in the set of acI genomes, and counted each COG as present in a clade if that COG was present in at least one genome belonging to that clade. We then used BBMap to map metatranscriptome reads to our reference genome collection, and calculated gene expression for each COG on a Reads Per Kilobase Million (RPKM) basis (Figure 2).

Sequencing of cDNA from all four rRNA-depleted metatranscriptome samples yielded approximately 160 million paired-end reads. After merging, filtering, and further *in-silico* rRNA removal, approximately 81 million, or 51% of the reads remained (Table S5). After mapping the metatranscriptomes to our acI genomes, we calculated the average coverage of each genome in our reference collection. Within each clade, the most abundant genome was detected with at least 16-fold coverage (Table S6).

We then examined the number of metatranscriptomic reads that mapped to the COGs identified within each clade. OrthoMCL identified a total of 5013 protein clusters across the three clades (Table S7) with an average confidence of 84% in annotation for COGs containing more than one gene. The COGs were unequally distributed across the three clades, with clade acI-A genomes containing 3175 COGs (63%), clade acI-B genomes containing 3459 COGs (69%), and clade acI-C genomes containing 1365 COGs (27%). Of these, 525 COGs were expressed in clade acI-A, 661 in clade acI-B, and 813 in clade acI-C (Table S8). Among expressed genes, the median log2 RPKM value was 31.1 in clade acI-A, 32.0 in clade acI-B, and 69.4 in clade acI-C.

### Auxotrophies and Nutrient Sources of the acI Lineage

Seed set analysis yielded seven auxotrophies that could be readily mapped to ecophysiological attributes of the acI lineage (Figure 3A and Table S3). In all three clades, beta-alanine was identified as a seed compound, suggesting an auxotrophy for pantothenic acid (Vitamin B5), a precursor to coenzyme A formed from beta-alanine and pantoate (Table S9). In bacteria, beta-alanine is typically synthesized via aspartate decarboxylation, and we were unable to identify a candidate gene for this enzyme (aspartate 1-decarboxylase, E.C. 4.1.1.11) in any acI genome. Pyridoxine 5’-phosphate and 5’-pyridoxamine phosphate (forms of the enzyme cofactor pyridoxal 5’-phosphate, Vitamin B6) were also predicted to be seed compounds, and numerous enzymes in the biosynthesis of these compounds were not found in the genomes (Table S9).

Clades within the acI lineage also exhibited distinct auxotrophies. Clade acI-A was predicted to be auxotrophic for the cofactor tetrahydrofolate (THF or Vitamin B9), and numerous enzymes for its biosynthesis were missing (Table S9). This cofactor plays an important role in the metabolism of amino acids and vitamins. In turn, clade acI-B was predicted to be auxotrophic for adenosylcobalamin (Vitamin B12), containing only four reactions from its biosynthetic pathway (Table S9). Finally, acI-C was predicted to be auxotrophic for the nucleotide uridine monophosphate (UMP, used as a monomer in RNA synthesis) and the amino acids lysine and homoserine. In all cases multiple enzymes for the biosynthesis of these compounds were not found in the acI-C genomes (Table S9).

A number of seed compounds were also predicted to be degraded by members of the acI lineage (Figure 3B and Table S3). Both clades acI-A and acI-B were predicted to use D-altronate and trans-4-hydroxy proline as nutrients, and acI-B was additionally predicted to use glycine betaine.

Finally, all three clades were predicted to use di-peptides and the sugar maltose as nutrients. Clades acI-A and acI-C were also predicted to consume the polysaccharides stachyose, manninotriose, and cellobiose. In all cases, these compounds were associated with reactions catalyzed by peptidases or glycoside hydrolases (Table S10 and S11), which may be capable of acting on compounds beyond the predicted seed compounds. Thus, we used these annotations to define nutrient sources, rather than using the predicted seed compounds themselves. Among these nutrient sources were di- and polypeptides, predicted to be released from both cytosolic- and membrane-bound aminopeptidases. As discussed below, we identified a number of transport proteins capable of transporting these released residues. In Lake Mendota, clades acI-B and acI-C expressed two aminopeptidases, one of which was expressed at upwards of 175% of the median gene expression levels (Table S10). Clade acI-A expressed a third aminopeptidase at a lower level of 40% the median gene expression level (Table S10).

All three clades were predicted to encode an alpha-glucosidase, which in Lake Mendota was only expressed in clades acI-B and acI-C, at upwards of 60% of the median gene expression level (Table S11). All three clades also encode a beta-glucosidase, but it was not expressed in our samples. Furthermore, all three clades encode an alpha-galactosidase and multiple maltodextrin glucosidases (which free maltose from maltotriose), but these were only expressed in clades acI-A and acI-C. The alpha-galactosidase had a log2 RPKM expression value of 1.5 times the median in clade acI-C, while the maltodextrin glucosidases were expressed at approximately 30% of the median (Table S11) in both clades acI-A and acI-C.

### Compounds Transported by the acI Lineage

Microbes may be capable of transporting compounds that are not strictly required for growth, and comparing such compounds to predicted seed compounds can provide additional information about an organism’s ecology. Thus, we used the metabolic network reconstructions for the acI genomes to systematically characterize the transport capabilities of the acI lineage.

All acI clades encode for and expressed a diverse array of transporters (Figure 4, Tables S12 and S13, and the Supplemental Text S1). Consistent with the presence of peptidases, all clades contain numerous genes for the transport of peptides and amino acids, including putative oligopeptide and branched-chain amino acid transporters, as well as putative transporters for the polyamines spermidine and putrescine. All clades also contain a putative transporter for ammonium. The ammonium, branched-chain amino acid, and oligopeptide transporters had expression values above the median, with expression values for the substrate-binding protein (of the ATP-binding cassette (ABC) transporters) ranging from 1.7 to 411 times the median (Table S13). In contrast, while all clades expressed some genes from the polyamine transporters, only clade acI-B expressed the binding protein, at approximately 27.8 times the median (Table S13). Finally, clades acI-A and acI-B also contain a putative transporter for glycine betaine, which was only expressed in clade acI-A, at approximately 9.6 times the median (Table S13). However, we cannot rule out the possibility that the expression of these transporters changes with space and time, and that all three clades may express these enzymes under a different condition.

**Figure 4.**
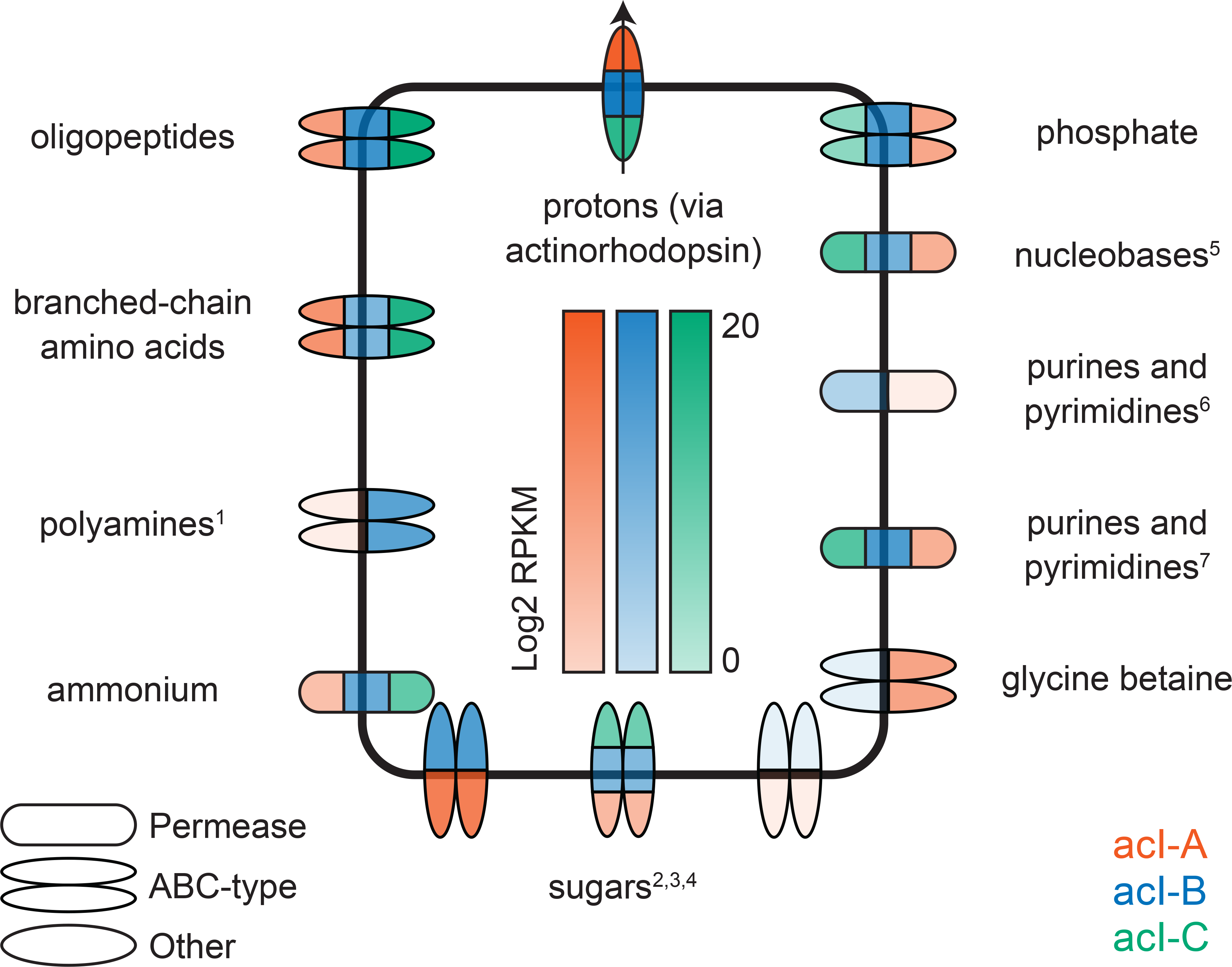
Transporters that are actively expressed by members of the acI lineage, as inferred from consensus annotations of genes associated with transport reactions present in metabolic network reconstructions. The intensity of the color indicates the log2 RPKM of the encoding gene cluster. For multi-subunit transporters, the RPKM of the substrate-binding subunit was chosen (Table S13). For some transporters, consensus annotations have been replaced with broad metabolite classes. Such metabolite classes are indicated with superscripts, and the original annotations are as follows: 1 - spermidine, putrescine; 2 - maltose; 3 - xylose; 4 -ribose; 5 - uracil; 6 cytosine / purine / uracil / thiamine / allantoin; 7 - xanthine / uracil / thiamine / ascorbate.

All clades also expressed transporters consistent with the presence of glycoside hydrolases, including transporters annotated as putative maltose, xylose, and ribose ABC-type transporters, which may indicate that acI bacteria are capable of transporting sugars, including both di- (maltose) and mono-saccharides (xylose and ribose). Of these, the putative maltose transporter was most highly expressed (but only in clades acI-A and acI-B), with expression values for the substrate-binding protein ranging in excess of 40 times the median (Table S13).

Representatives from the acI lineage also encode and expressed a number of transporters that do not have corresponding seed compounds, including potential nucleobase and purine/pyrimidine transporters (annotated as a uracil and a xanthine/uracil/thiamine/ascorbate family permease, respectively). Both of these are expressed in all three clades, with expression values ranging from 4.7 to 46 times the median (Table S13). Clades acI-A and acI-B also contain a second potential purine/pyrimidine transporter (annotated as a cytosine/purine/uracil/thiamine/allantoin family permease), which was only expressed in clade acI-B (Table S13). These transporters may be responsible for the uptake of the seed compounds UMP (a pyrimidine derivative) and Vitamin B1 (also known as thiamine). In addition, clade acI-A contains but did not express a putative transporter for cobalamin (Vitamin B12), and both clades acI-A and acI-B contain but did not express transporters for thiamin (Vitamin B1) and biotin (Vitamin B7) (Table S13).

Finally, all three clades expressed actinorhodopsin, a light-sensitive protein that is expected to function as a proton efflux pump (30). In all clades, actinorhodopsin was among the top ten most highly-expressed genes (Table S7), with expression values in excess of 84 times the median in all three clades (Table S7). Given that many of the transport proteins are ABC transporters, we speculate that actinorhodopsin may facilitate maintenance of the proton gradient necessary for ATP synthesis. Coupled with high expression levels of diverse transporters, this result strongly suggests that acI functions as a photoheterotroph. However, it remains to be seen if this behavior is a general feature of acI physiology or restricted to the specific conditions of the lake and our sampling period.

## Discussion

This study uses high-throughput metabolic network reconstruction and the seed set framework to predict auxotrophies and nutrient sources of uncultivated microorganisms from incomplete genome sequences. The computational approach easily scales to hundreds of metabolic reconstructions, and enables a targeted analysis by identifying those compounds through which an organism interacts with its environment. However, predicted seed compounds are sensitive to the metabolic network structure, and analyzing the results requires significant manual curation of the metabolic reconstruction and accurate interpretation of the underlying gene annotations. As a consequence, the seed set framework as not as high-throughput as initially envisioned, but is nevertheless suitable for analysis of microorganisms with high-quality metabolic network reconstructions.

Our predictions of substrate use capabilities of the acI lineage are largely congruent with previous genome-centric studies based on smaller but manually curated genome collections (22, 25, 27), indicating that the use of automatic metabolic network reconstructions yields similar predictions to comparatively more manual metabolic reconstruction efforts, while being both more high-throughput and focused on an organism’s substrate utilization capabilities. In particular, this study predicts that the consumption of N-rich compounds is a universal feature of the acI lineage, with all three clades predicted to consume ammonium, branched-chain amino acids, polyamines, and di- and oligopeptides. These findings agree with MAR-FISH studies that confirm the ability of acI bacteria to consume a variety of amino acids (20, 23). Furthermore, the presence of alpha- and beta-glucosidases are consistent with observations that acI bacteria consume glucose (19, 23), even though no obvious glucose transport system was found in the genomes. Because transport proteins are often capable of acting on multiple substrates, one of the putative sugar transporters may be responsible for glucose uptake activity.

However, our approach failed to recapitulate other genomic and experimental observations, including the uptake of N-acetylglucosamine (NAG) (31–33), the deoxynucleoside thymidine (23, 34), and acetate (19), and the potential to hydrolyze the cyanobacterial peptide cyanophycin via the enzyme cyanophycinase (22, 25). Inspection of these discrepancies reveals some important limitations of the seed set framework and automatic metabolic reconstructions. First, the seed set framework only identifies compounds that the metabolic network must obtain from its environment, and will fail to identify compounds that the organism can acquire from its environment but can also synthesize itself. Thymidine and acetate fall into this category. Second, automatic metabolic network reconstructions may not fully capture an organism’s metabolic network (e.g., due to missing or incorrect genome annotations). Manual inspection of the previously-identified cyanophycinase gene revealed that KBase annotated this putative enzyme as a hypothetical protein. As biochemical characterization of hypothetical proteins and automatic gene and protein annotation are active areas of research, we anticipate that advances in these fields will continue to improve the accuracy of automatic metabolic network reconstructions.

This study also suggests that auxotrophies for some vitamins may be universal features of the acI lineage, as we predict all clades to be auxotrophic for pantothenic acid and pyridoxal 5’-phosphate (Vitamins B5 and B6). We also predict new auxotrophies within the acI lineage, including THF (clade acI-A), adenosylcobalamin (Vitamin B12, clade acI-B), and lysine, homoserine, and UMP (clade acI-C). However, with the exception of adenosylcobalamin, we did not identify transporters for any of these compounds. This negative result may reflect our limited knowledge of transport proteins (35): transporters for these compounds may yet be present in the genomes, or one or more of the predicted transporters may act on these compounds. Furthermore, because the acI-C composite genome was estimated to be around 75% complete, we cannot rule out the possibility that the missing genes might be found in this clade when additional genomes are recovered. Nonetheless, these results provide additional support to the hypothesis that distributed metabolic pathways and metabolic complementarity may be common features of freshwater bacterial communities (36, 37).

Combined, these results suggest that acI are photoheterotrophs, making a living on a diverse array of N-rich compounds, saccharides, and light. The acI lineage does not appear to be metabolically self-sufficient, and may participate in the turnover of high molecular weight dissolved organic compounds, such as starch, glycogen, and cellulose. Metatranscriptomic analysis showed that transport proteins were among the most highly expressed in the acI genomes, and the expression of multiple putative amino acid transporters may facilitate uptake of these labile compounds. We also observed differences in the relative expression of these transporters, which may point to clade-specific differences in the affinity for these substrates. Finally, the actinorhodopsin protein was highly expressed, and may facilitate synthesis of the ATP needed to drive acI’s many ABC-type transporters.

Finally, the fragmented and incomplete nature of SAGs and MAGs required us to construct composite genomes for individual acI clades by leveraging multiple genomes from closely related populations. Such an approach limits the resolution of predictions, as we cannot make predictions at the level of tribes, smaller populations, or individual cells. Thus, metabolic diversification at these taxonomic levels will be missed. Constructing composite genomes may also overestimate the metabolic capabilities of a clade or group: for example, if a complete pathway is present in a clade but distributed among different tribes, the clade will only be able to carry out the entire pathway *in situ* if all tribes are present in close enough proximity to exchange pathway intermediates. Nonetheless, the seed set approach provides a framework that can be used to generate new hypotheses about the substrates used by members of a defined phylogenetic group, provided multiple closely related genomes are available. As metagenomic assembly and binning techniques and single cell sequencing methods improve and complete genomes become available, we anticipate our approach being applied to individual microbial genomes.

## Materials and Methods

### A Freshwater Reference Genome Collection

This study relies on an extensive collection of freshwater bacterial genomes, containing MAGs obtained from two metagenomic time-series from two Wisconsin lakes (27, 38), as well as SAGs from three lakes in the United States (21). Additional information about this genome collection can be found in the Supplemental Text S1.

### Metatranscriptome Sampling and Sequencing

This study used four metatranscriptomes obtained as part of a larger study of gene expression in freshwater microbial communities. Additional information about these samples can be found in the Supplemental Text S1. All protocols and scripts for sample collection, RNA extraction, rRNA depletion, sequencing, and bioinformatic analysis can be found on Github (https://github.com/McMahonLab/OMD-TOIL, DOI:######). Metadata for the four samples used in this study can be found in Table S6, and the raw RNA sequences can be found on the National Center for Biotechnology Information’s Sequence Read Archive (SRA) under BioProject PRJNA362825.

### Identification of acI SAGs and Actinobacterial MAGs

The acI were previously phylogenetically divided into three clades (acI-A, acI-B, and acI-C) and thirteen tribes on the basis of their 16S rRNA gene sequences (28). The acI SAGs were identified within a previously-published genome collection (21) and classified to the tribe level using partial 16S rRNA genes and a reference taxonomy for freshwater bacteria, as described in the Supplemental Text S1. Actinobacterial MAGs were identified within two metagenomic time-series (27, 38) using taxonomic assignments from a subset of conserved marker genes, as described in the Supplemental Text S1. Phylogenetic analysis of acI SAGs and Actinobacterial MAGs was performed using a concatenated alignment of single-copy marker genes obtained via Phylosift (39). Maximum likelihood trees were generated using RAxML (40) using the automatic protein model assignment option (PROTGAMMAAUTO) and 100 bootstraps.

### Genome Annotation, Metabolic Network Reconstruction, and Computation and Evaluation of Seed Compounds

In the seed set framework, an organism’s metabolism is represented via a metabolic network graph, in which nodes denote compounds and edges denote enzymatically-encoded biochemical reactions linking substrates and products (41). Allowable biochemical transformations can be identified by drawing paths along the network, in which a sequence of edges connects a sequence of distinct vertices. In our implementation of the seed set framework, metabolic network graphs were generated as follows.

Genome annotations were performed and metabolic network reconstructions were built using KBase. Contigs for each genome were uploaded to KBase and annotated using the “Annotate Microbial Contigs” method with default options, which uses components of the RAST toolkit for genome annotation (42, 43). Metabolic network reconstructions were obtained using the “Build Metabolic Model” app with default parameters, which relies on the Model SEED framework (44) to build a draft metabolic model. No gap-filling was performed, to ensure that the reconstructions only contained reactions with genomic evidence. These reconstructions were then pruned (currency metabolites and highly-connected compounds) and converted to metabolic network graphs (Figure S3 and Supplemental Text S1). Many of the individual acI genomes are incomplete. Therefore, composite metabolic network graphs were constructed for each tribe and clade, to increase the accuracy of seed identification by means of a more complete metabolic network (Figure S4 and Supplemental Text S1).

Formally, the seed set of the network is defined as the minimal set of compounds that cannot be synthesized from other compounds in the network, and whose presence enables the synthesis of all other compounds in the network (1). Seed compounds for each composite metabolic network graph were calculated using a new Python implementation of the seed set framework (1) (Figure S5 and the Supplemental Text S1). Because seed compounds are computed from a metabolic network, it is important to manually evaluate all predicted seed compounds to identify those that may be biologically meaningful, and do not arise from errors in the metabolic network reconstruction. Compounds involved in fatty acid and phospholipid biosynthesis pathways were removed during curation, as these pathways are often organism-specific and unlikely to be properly annotated by automatic metabolic reconstruction pipelines. Seed compounds related to currency metabolites (compounds used to carry electrons and functional groups) were also removed, as reactions for the synthesis of these compounds may have been removed during network pruning.

The Supplemental Text S1 contains a series of brief vignettes explaining why select compounds were discarded based on the afore-mentioned considerations, and provides examples of additional curation efforts applied to biologically plausible compounds. For a plausible auxotrophy, we screened the genomes for the canonical biosynthetic pathway(s) for that compound, and retained those compounds for which the biosynthetic pathway was incomplete. For a plausible nutrient source, we screened the genomes for the canonical degradation pathway(s) for that compound, and retained those compounds for which the degradation pathway was complete.

All computational steps were implemented using Python scripts, freely available as part of the reverseEcology Python package developed for this project (https://pypi.python.org/pypi/reverseEcology/, DOI:######).

### Identification of Transported Compounds

For each genome, we identified all transport reactions present in its metabolic network reconstruction. Gene-protein-reaction associations (GPRs) for these reactions were manually curated to remove unannotated proteins, group genes into operons (if applicable), and to identify missing subunits for multi-subunit transporters. These genes were then mapped to their corresponding COGs, and grouped accordingly. Finally, the most common annotation for each COG was used to identify likely substrates for each of these groups. Only transporters with >50% confidence in the substrate-binding subunit were retained. Because identification and annotation of transport proteins is an active area of research (35), substrates for each transporter are described as putative and acting on molecular classes (e.g., saccharide, amino acid) instead of specific compounds, to better reflect the promiscuity of transport proteins and the ambiguity of their annotation.

### Protein Clustering, Metatranscriptomic Mapping, and Clade-Level Gene Expression

OrthoMCL (29) was used to identify clusters of orthologous groups (COGs) in the set of acI genomes. Both OrthoMCL and BLAST were run using default options (45). Annotations were assigned to protein clusters by choosing the most common annotation among all genes assigned to the respective cluster and a confidence score assigned to each COG (fraction of genes having the most common annotation). Trimmed and merged metatranscriptomic reads from each of the four biological samples were then pooled and mapped to a single reference fasta file containing all acI genomes using BBMap (https://sourceforge.net/projects/bbmap/) with the *ambig=random* and *minid=0.95* options. The 95% identity cutoff was chosen as this represents a well-established criterion for identifying microbial species using average nucleotide identity (ANI) (46), while combining the *ambig* option with competitive mapping using pooled acI genomes as the reference ensures that reads map only to a single genome. These results were then used to compute the expression of each COG in each clade.

Next, HTSeq-Count (47) was run using the *intersection_strict* option to count the total number of reads that map to each gene in our acI genome collection. After mapping, the list of counts was filtered to remove those genes that did not recruit at least ten reads. Using the COGs identified by OrthoMCL, the genes that correspond to each COG were then identified.

Within each clade, gene expression for each COG was computed on a Reads Per Kilobase Million (RPKM) basis (48), while also accounting for different gene lengths within a COG and numbers of mapped reads for each genome within a clade. That is, the RPKM value for a single COG represents the sum of RPKM values for each gene within that COG, normalized to the appropriate gene length and total number of mapped reads. RPKM counts were then normalized to the median level of gene expression within that clade. Finally, the expression data (mapping of transcript reads to genes) were visualized to ensure RPKM calculations were based on continuous transcription of each gene.

### Availability of Data and Materials

All genomic and metatranscriptomic sequences are available through IMG and NCBI, respectively. A reproducible version of this manuscript is available at https://github.com/joshamilton/Hamilton_acI_2016 (DOI:######).

## Acknowledgements

We thank past and present members of the McMahon lab for collecting water samples for DNA and RNA sequencing and Frank Aylward for guidance on metatranscriptomic analysis. DNA sequencing was supported through the JGI Community Science Program. The work conducted by the JGI, a DOE Office of Science User Facility, is supported by the Office of Science of the U.S. Department of Energy under Contract No. DE-AC02-05CH11231. This material is based upon work that is supported by the National Institute of Food and Agriculture, U.S. Department of Agriculture, under award number 2016-67012-24709 to JJH and WIS01789 to KDM. KDM also acknowledges funding from the United States National Science Foundation (NSF) Microbial Observatories program (MCB-0702395), the NSF Long Term Ecological Research program (NTL-LTER DEB-1440297), an NSF INSPIRE award (DEB-1344254), and the University of Wisconsin System. KDM and KTF acknowledge National Oceanic and Atmospheric Administration (NOAA) grant #NA10OAR4170070, Wisconsin Sea Grant College Program Project #HCE-25, through NOAA’S National Sea Grant College Program, U.S. Department of Commerce. The funders had no role in study design, data collection and interpretation, or the decision to submit the work for publication.

---

**Supplementary Figure 1.**
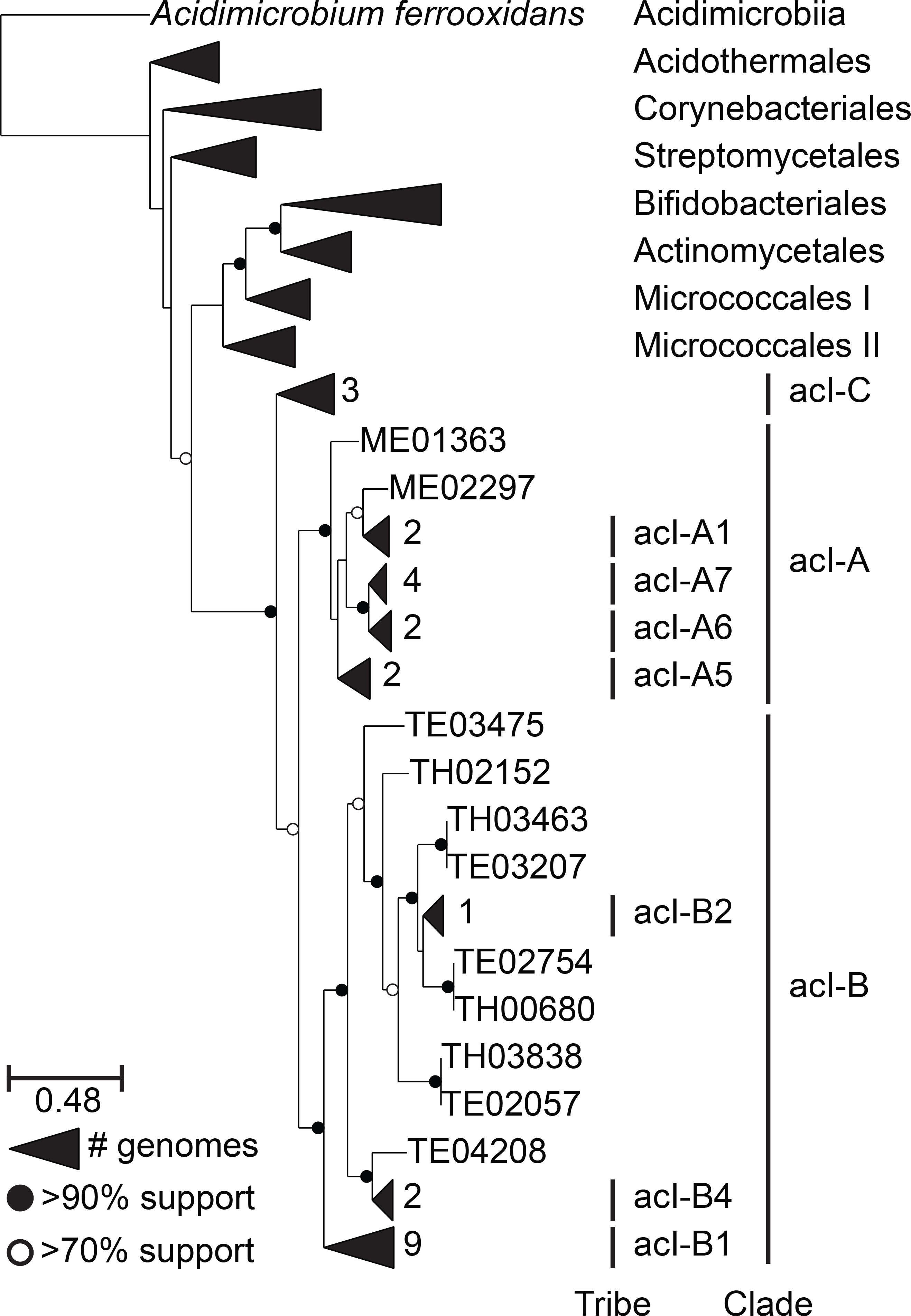
Phylogenetic placement of the genomes used in this study within the acI lineage, relative to other sequenced actinobacterial genomes in the class Actinobacteria (49) (Table S19). The tree was built using RAxML (40) from a concatenated alignment of protein sequences from 37 single-copy marker genes (39). The class Acidimicrobiia forms the outgroup. Vertical black bars indicate groups of genomes belonging to defined tribes/clades within the acI lineage, as determined using 16S rRNA gene sequences (for SAGs and bin FNEF8-2 bin_7 acI-B only) and a defined taxonomy (28). SAGs are indicated with italic text.

**Supplementary Figure 2.**
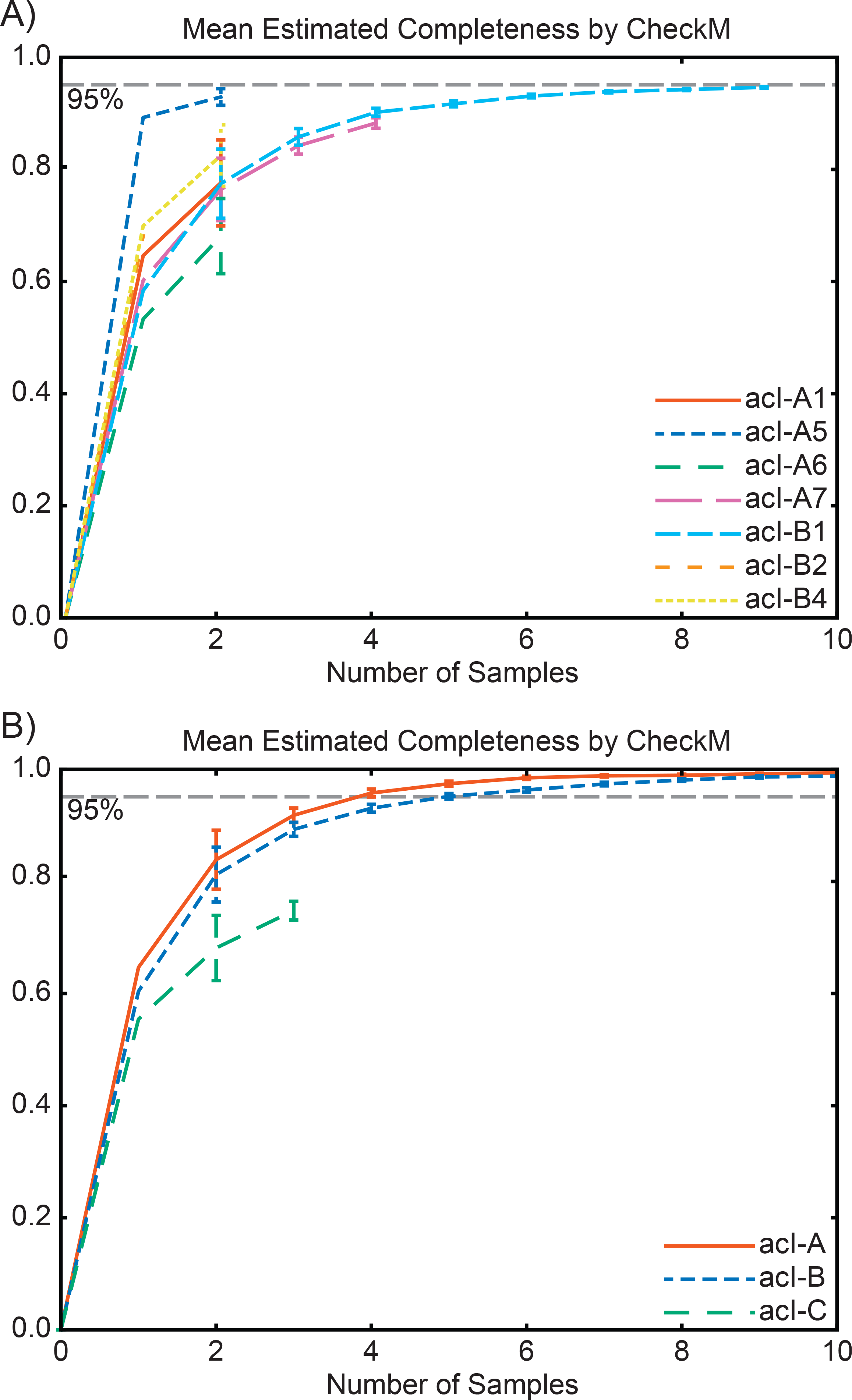
Mean estimated completeness of tribe-level (clade-level) population genomes as a function of the number of sampled genomes. For each tribe (clade), genomes were randomly sampled (with replacement) from the set of all genomes belonging to that tribe (clade). Completeness was estimated using 204 single-copy marker genes from the phylum Actinobacteria (50). Error bars represent the 95% confidence interval estimated from 1000 iterations.

**Supplementary Figure 3.**
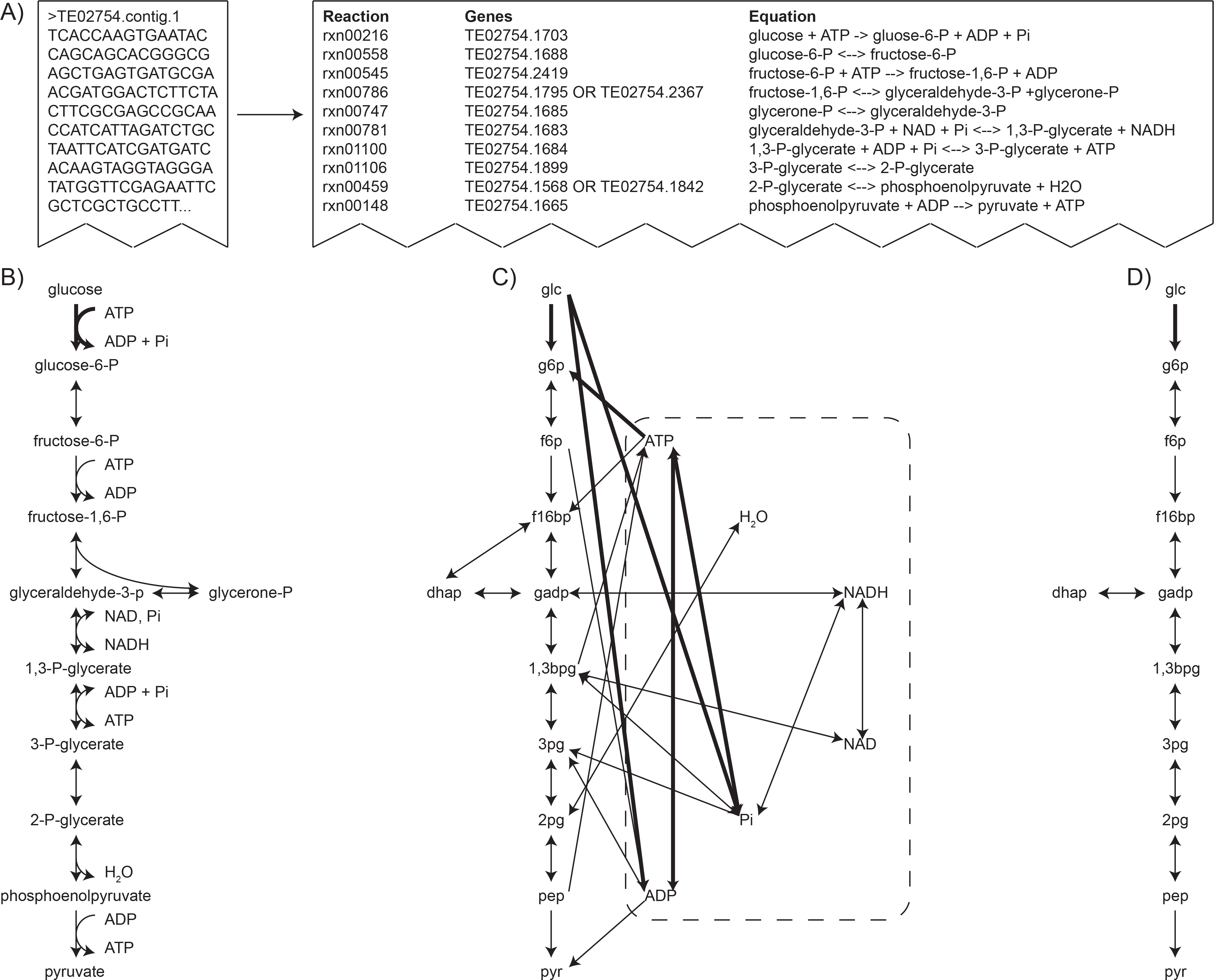
Converting an unannotated genome to a metabolic network graph, for a simplified genome containing only glycolysis. **(A)** Microbial contigs are annotated using KBase, and a metabolic network reconstruction is built from the annotations. The reconstruction provides links between protein-encoding genes in the genome and the enzymatic reactions catalyzed by those proteins. **(B)** The metabolic network reconstruction represents metabolism as a hypergraph, in which metabolites are represented as nodes and reactions as hyperedges. In this representation, an edge can connect more than two nodes. For example, a single hyperedge (denoted by a heavy black line) connects the metabolites glucose and ATP to glucose-6P, ADP, and Pi. For clarity, protons are not shown. **(C)** However, the algorithm used by the seed set framework requires metabolism to be represented as a metabolic network graph, in which an edge can connect only two nodes. In this representation, a reaction is represented by a set of edges connecting all substrates to all products. For example, the heavy hyperedge in (B) is now denoted by six separate edges connecting glucose to ADP, glucose to Pi, glucose to glucose-6P, ATP to ADP, ATP to Pi, and ATP to glucose-6P (again denoted by heavy black lines). Of these, only one (glucose to glucose-6P) is biologically meaningful. The dotted line surrounds the currency metabolites. **(D)** The metabolic network graph is then pruned, a process which removes all currency metabolites and any edges in which those metabolites participate. Of the six heavy edges in (C), only the biologically meaningful one is retained, connecting glucose to glucose-6P (again denoted by a heavy black line). The images in (B) and (C) are modified from (51).

**Supplementary Figure 4.**
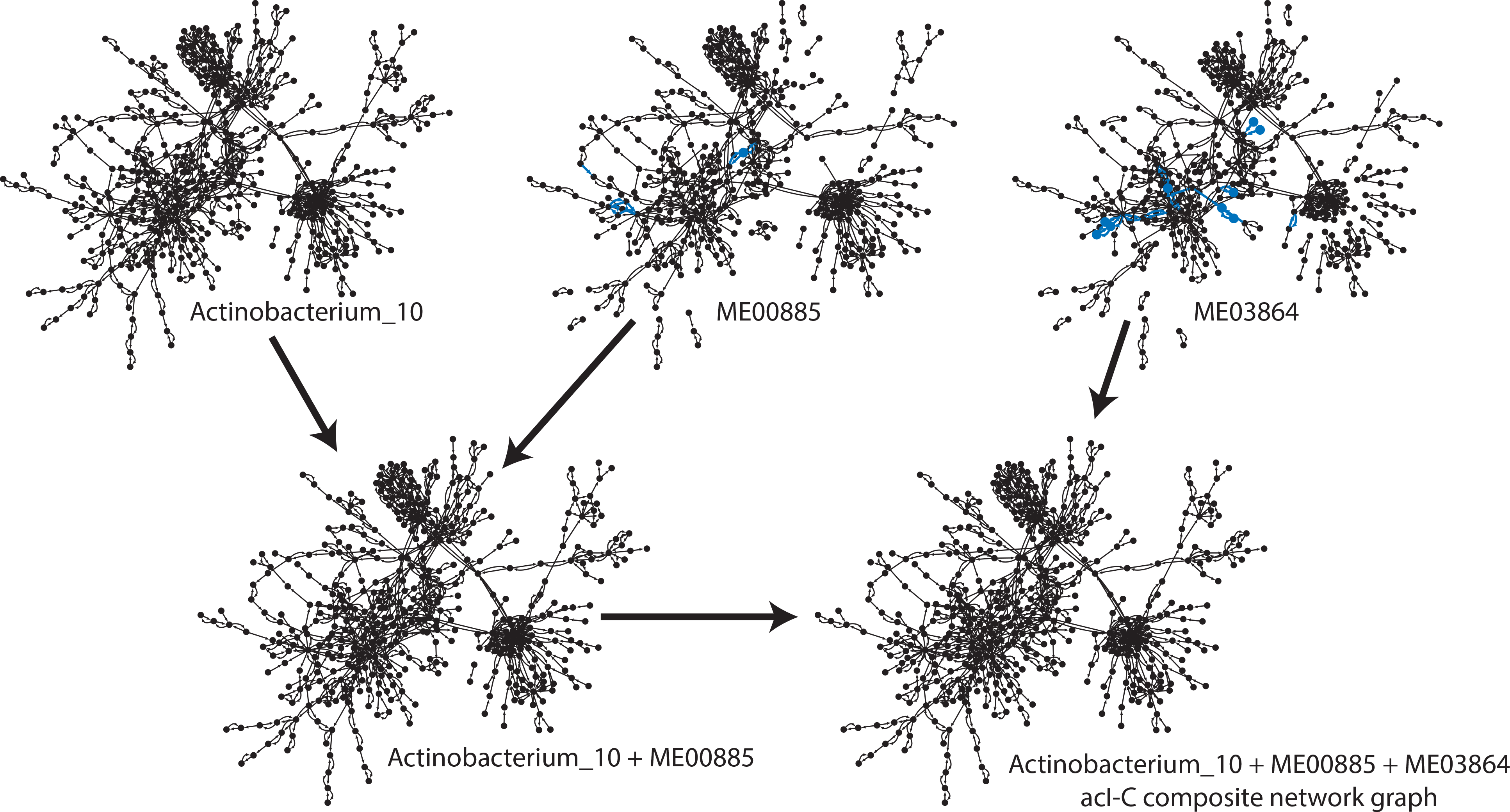
Construction of composite metabolic network graph for clade acI-C. Beginning with metabolic network graphs for genomes Actinobacterium_10 and ME00885, nodes and edges unique to ME00885 are identified (in blue). These nodes and edges are added to the Actinobacterium_10 graph, giving the composite metabolic network graph for these two genomes (Actinobacterium_10 + ME00885). Then, this graph is compared to the graph for ME03864, and nodes and edges unique to ME03864 are identified (in blue). These nodes and edges are added to the Actinobacterium_10 + ME00885 metabolic network graph, giving the composite metabolic network graph for clade acI-C.

**Supplementary Figure 5.**
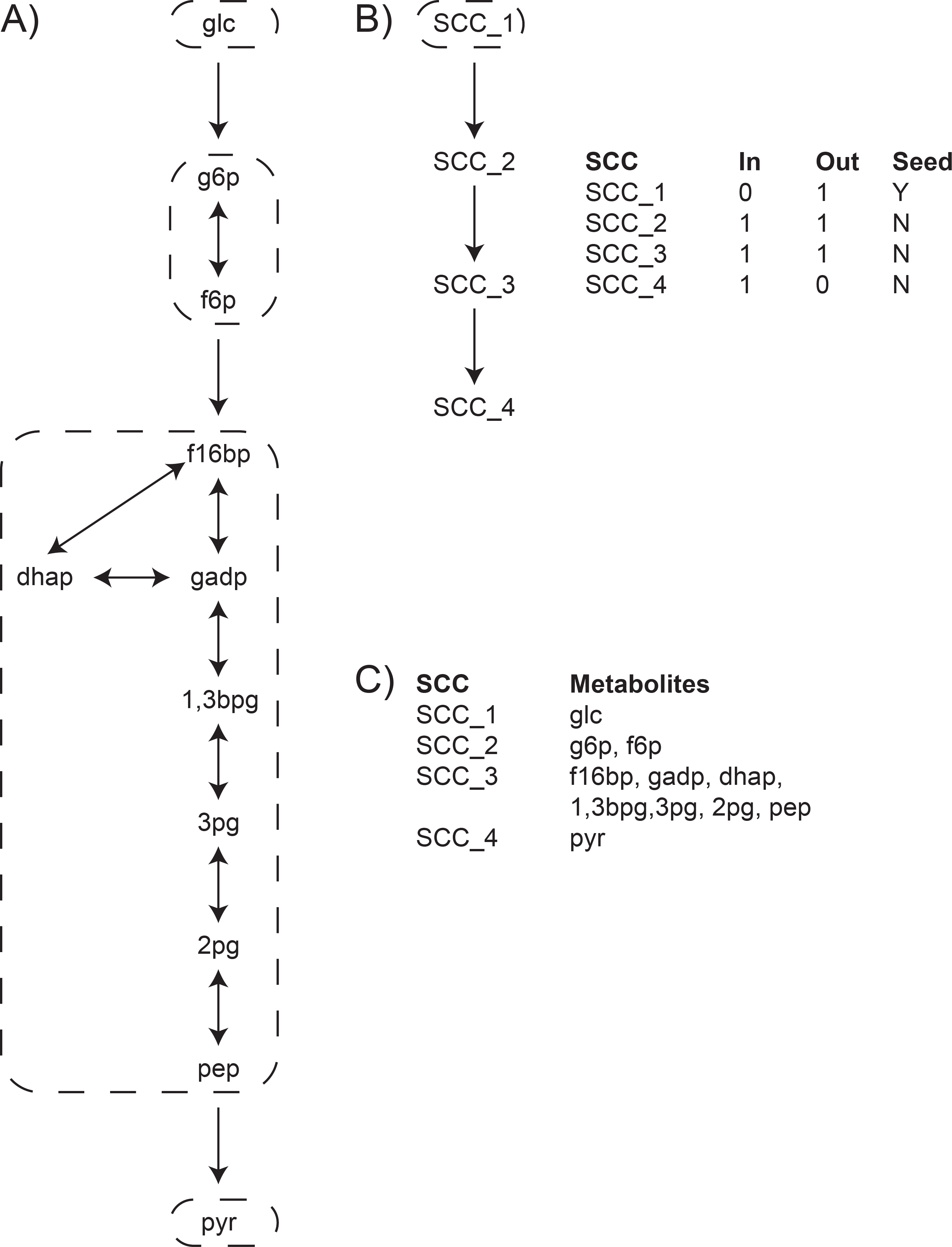
Identifying seed compounds in metabolic networks, using the same metabolic network as in Supplementary Figure S1. **(A)** To identify seed compounds, the metabolic network graph is first decomposed into its strongly connected components (SCCs), sets of nodes such that each node in the set is reachable from every other node. Here, each set of circled nodes corresponds to a unique SCC. **(B)** SCC decomposition enables seed sets to be identified from source components (components with no incoming edges) on the condensation of the original graph. In the condensation of the original graph shown here, each node corresponds to a unique SCC. This network has a single seed set, SCC_1, enclosed in a dotted circle. **(C)** Seed compounds can be found from the mapping between SCCs and their constituent metabolites. In this example, glucose is the sole seed compound. While this particular result is probably intuitive, real metabolic networks are considerably more complex. Note: The visual representations shown here are intended to illustrate the metabolic network reconstruction process, and are not indicative of the data structures used by our pipeline.

**Supplementary Figure 6.**
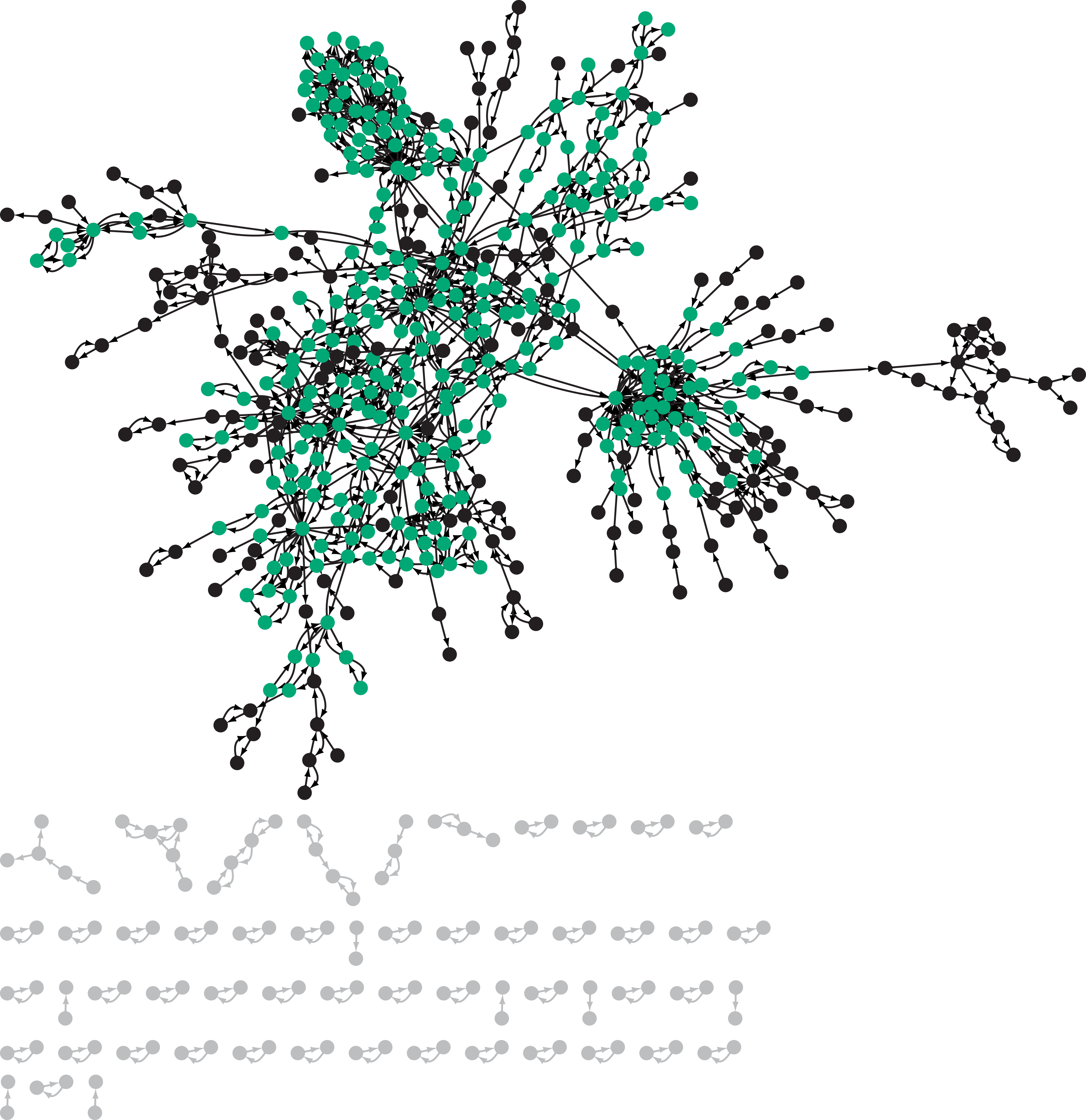
Complete composite metabolic network graph for clade acI-C, showing disconnected components (gray) and the single largest component (green and black). Disconnected components are dropped prior to computing the network’s seed sets because these groups of nodes are not connected to the bulk of the network. Within the single largest component, the giant strong component contains a substantial fraction of the compounds (green nodes), giving rise to a bow-tie structure in the metabolic network graph.

Supplementary Tables S1 to S19.

## References

1. Borenstein E, Kupiec M, Feldman MW, Ruppin E. 2008. Large-scale reconstruction and phylogenetic analysis of metabolic environments. Proceedings of the National Academy of Sciences 105:14482–14487.

2. Falkowski PG, Fenchel T, Delong EF. 2008. The microbial engines that drive Earth’s biogeochemical cycles. Science 320:1034–1039.

3. Blaser MJ, Cardon ZG, Cho MK, Dangl JL, Donohue TJ, Green JL, Knight R, Editor S, Maxon ME, Northen TR, Pollard KS, Editor S. 2016. Toward a Predictive Understanding of Earth’s Microbiomes to Address 21st Century Challenges. mBio 7:e00074–16.

4. Sangwan N, Xia F, Gilbert JA. 2016. Recovering complete and draft population genomes from metagenome datasets. Microbiome 4:8.

5. Lawson CE, Wu S, Bhattacharjee AS, Hamilton JJ, Mcmahon KD, Goel R, Noguera DR. 2017. Metabolic network analysis reveals microbial community interactions in anammox granules. Nature Communications 8:15416.

6. Anantharaman K, Brown CT, Hug LA, Sharon I, Castelle CJ, Probst AJ, Thomas BC, Singh A, Wilkins MJ, Karaoz U, Brodie EL, Williams KH, Hubbard SS, Banfield JF. 2016. Thousands of microbial genomes shed light on interconnected biogeochemical processes in an aquifer system. Nature Communications 7:13219.

7. Orth JD, Thiele I, Palsson BØ. 2010. What is flux balance analysis? Nature Biotechnology 28:245–248.

8. Arkin AP, Stevens RL, Cottingham RW, Maslov S, Henry CS, Dehal P, Ware D, Perez F, Harris NL, Canon S, Sneddon MW, Henderson ML, Riehl WJ, Gunter D, Mur D, Yoo S. 2016. The DOE Systems Biology Knowledgebase (KBase). bioRxiv.

9. Zwart G, Hiorns WD, Methé BA, Agterveld van MP, Huismans R, Nold SC, Zehr JP, Laanbroek HJ. 1998. Nearly Identical 16S rRNA Sequences Recovered from Lakes in North America and Europe Indicate the Existence of Clades of Globally Distributed Freshwater Bacteria. Systematic and Applied Microbiology 21:546–556.

10. Glockner FO, Zaichikov E, Belkova N, Denissova L, Pernthaler J, Pernthaler A, Amann R. 2000. Comparative 16S rRNA analysis of lake bacterioplankton reveals globally distributed phylogenetic clusters including an abundant group of actinobacteria. Applied and Environmental Microbiology 66:5053–5065.

11. Zwart G, Crump BC, Kamst-van Agterveld MP, Hagen F, Han SK. 2002. Typical freshwater bacteria: An analysis of available 16S rRNA gene sequences from plankton of lakes and rivers. Aquatic Microbial Ecology 28:141–155.

12. Newton RJ, Kent AD, Triplett EW, McMahon KD. 2006. Microbial community dynamics in a humic lake: Differential persistence of common freshwater phylotypes. Environmental Microbiology 8:956–970.

13. Wu QL, Zwart G, Schauer M, Kamst-Van Agterveld MP, Hahn MW. 2006. Bacterioplankton community composition along a salinity gradient of sixteen high-mountain lakes located on the Tibetan Plateau, China. Applied and Environmental Microbiology 72:5478–5485.

14. Newton RJ Jones SE Helmus MR McMahon KD. 2007. Phylogenetic ecology of the freshwater Actinobacteria acI lineage. Applied and Environmental Microbiology 73:7169–7176.

15. Wu X Xi W Ye W Yang H. 2007. Bacterial community composition of a shallow hypertrophic freshwater lake in China revealed by 16S rRNA gene sequences. FEMS Microbiology Ecology 61:85–96.

16. De Wever A Van Der Gucht K Muylaert K Cousin S Vyverman W. 2008. Clone library analysis reveals an unusual composition and strong habitat partitioning of pelagic bacterial communities in Lake Tanganyika. Aquatic Microbial Ecology 50:113–122.

17. Humbert JF Dorigo U Cecchi P Le Berre B Debroas D Bouvy M. 2009. Comparison of the structure and composition of bacterial communities from temperate and tropical freshwater ecosystems. Environmental Microbiology 11:2339–2350.

18. Ghai R McMahon KD Rodriguez-Valera F. 2012. Breaking a paradigm: cosmopolitan and abundant freshwater actinobacteria are low GC. Environmental Microbiology Reports 4:29–35.

19. Buck U Grossart H-P, Amann RI Pernthaler J. 2009. Substrate incorporation patterns of bacterioplankton populations in stratified and mixed waters of a humic lake. Environmental Microbiology 11:1854–1865.

20. Salcher MM Pernthaler J Posch T. 2010. Spatiotemporal distribution and activity patterns of bacteria from three phylogenetic groups in an oligomesotrophic lake. Limnology and Oceanography 55:846–856.

21. Martinez-Garcia M Swan BK Poulton NJ Gomez ML Masland D Sieracki ME Stepanauskas R. 2012. High-throughput single-cell sequencing identifies photoheterotrophs and chemoautotrophs in freshwater bacterioplankton. The ISME Journal 6:113–123.

22. Garcia SL McMahon KD Martinez-Garcia M Srivastava A Sczyrba A Stepanauskas R Grossart H-P, Woyke T Warnecke F. 2013. Metabolic potential of a single cell belonging to one of the most abundant lineages in freshwater bacterioplankton. The ISME Journal 7:137–147.

23. Salcher MM Posch T Pernthaler J. 2013. In situ substrate preferences of abundant bacterioplankton populations in a prealpine freshwater lake. The ISME Journal 7:896–907.

24. Ghai R Mizuno CM Picazo A Camacho A Rodriguez-Valera F. 2014. Key roles for freshwater Actinobacteria revealed by deep metagenomic sequencing. Molecular Ecology 23:6073–6090.

25. Ghylin TW Garcia SL Moya F Oyserman BO Schwientek P Forest KT Mutschler J Dwulit-Smith J Chan L-K, Martinez-Garcia M Sczyrba A Stepanauskas R Grossart H-P, Woyke T Warnecke F Malmstrom RR Bertilsson S McMahon KD. 2014. Comparative single-cell genomics reveals potential ecological niches for the freshwater acI Actinobacteria lineage. The ISME Journal 8:2503–2516.

26. Tsementzi D Poretsky RS Rodriguez-R LM Luo C Konstantinidis KT. 2014. Evaluation of metatranscriptomic protocols and application to the study of freshwater microbial communities. Environmental Microbiology Reports 6:640–655.

27. Garcia SL Buck M McMahon KD Grossart H-P, Eiler A Warnecke F. 2015. Auxotrophy and intra-population complementary in the ‘interactome’ of a cultivated freshwater model community. Molecular Ecology 24:4449–4459.

28. Newton RJ Jones SE Eiler A McMahon KD Bertilsson S. 2011. A guide to the natural history of freshwater lake bacteria. Microbiology and Molecular Biology Reviews 75:14–49.

29. Li L Stoeckert CJ Roos DS. 2003. OrthoMCL: identification of ortholog groups for eukaryotic genomes. Genome Research 13:2178–89.

30. Sharma AK Zhaxybayeva O Papke RT Doolittle WF. 2008. Actinorhodopsins: Proteorhodopsin-like gene sequences found predominantly in non-marine environments. Environmental Microbiology 10:1039–1056.

31. Beier S Bertilsson S. 2011. Uncoupling of chitinase activity and uptake of hydrolysis products in freshwater bacterioplankton. Limnology and Oceanography 56:1179–1188.

32. Eckert EM Salcher MM Posch T Eugster B Pernthaler J. 2012. Rapid successions affect microbial N-acetyl-glucosamine uptake patterns during a lacustrine spring phytoplankton bloom. Environmental Microbiology 14:794–806.

33. Eckert EM Baumgartner M Huber IM Pernthaler J. 2013. Grazing resistant freshwater bacteria profit from chitin and cell-wall-derived organic carbon. Environmental Microbiology 15:2019–2030.

34. Pérez MT Hörtnagl P Sommaruga R. 2010. Contrasting ability to take up leucine and thymidine among freshwater bacterial groups: Implications for bacterial production measurements. Environmental Microbiology 12:74–82.

35. Saier MH Reddy VS Tamang DG Västermark Å. 2014. The transporter classification database. Nucleic Acids Research 42:D251–D258.

36. Garcia SL. 2016. Mixed cultures as model communities: hunting for ubiquitous microorganisms their partners and interactions. Aquatic Microbial Ecology 77:79–85.

37. Garcia SL Buck M Hamilton JJ Wurzbacher C Rosenblad MA McMaho KD Grossart H-P, Warnecke F Eiler A. 2017. Model communities hint to promiscuous metabolic linkages between ubiquitous free-living freshwater bacteria. bioRxiv.

38. Bendall ML Stevens SLR Chan L-K, Malfatti S Schwientek P Tremblay J Schackwitz W Martin J Pati A Bushnell B Froula J Kang D Tringe SG Bertilsson S Moran MA Shade AL Newton RJ McMahon KD Malmstrom RR. 2016. Genome-wide selective sweeps and gene-specific sweeps in natural bacterial populations. The ISME Journal 10:1589–1601.

39. Darling AE Jospin G Lowe E Matsen FA Bik HM Eisen JA. 2014. PhyloSift: phylogenetic analysis of genomes and metagenomes. PeerJ 2:e243.

40. Stamatakis A. 2014. RAxML version 8: a tool for phylogenetic analysis and postanalysis of large phylogenies. Bioinformatics 30:1312–1313.

41. Jeong H Tombor B Albert R Oltvai ZN Barabási A-L, Database I. 2000. The largescale organization of metabolic networks. Nature 407:651–654.

42. Brettin T Davis JJ Disz T Edwards RA Gerdes S Olsen GJ Olson R Overbeek RA Parrello B Pusch GD Shukla M Thomason JA Stevens R Vonstein V Wattam AR Xia F. 2015. RASTtk: a modular and extensible implementation of the RAST algorithm for building custom annotation pipelines and annotating batches of genomes. Scientific Reports 5:8365.

43. Overbeek RA Olson R Pusch GD Olsen GJ Davis JJ Disz T Edwards RA Gerdes SY Parrello B Shukla M Vonstein V Wattam AR Xia F Stevens RL. 2014. The SEED and the Rapid Annotation of microbial genomes using Subsystems Technology (RAST). Nucleic Acids Research 42:206–214.

44. Henry CS DeJongh M Best AA Frybarger PM Linsay B Stevens RL. 2010. Highthroughput generation optimization and analysis of genome-scale metabolic models. Nature Biotechnology 28:977–982.

45. Fischer S Brunk BP Chen F Gao X Harb OS Iodice JB Shanmugam D Roos DS Stoeckert CJ. 2011. Using OrthoMCL to assign proteins to OrthoMCL-DB groups or to cluster proteomes into new ortholog groups. Current Protocols in Bioinformatics Supplement:6.12.1.6–12.19.

46. Konstantinidis KT Tiedje JM. 2005. Genomic insights that advance the species definition for prokaryotes. Proceedings of the National Academy of Sciences 102:2567–2572.

47. Anders S Pyl PT Huber W. 2014. HTSeq A Python framework to work with highthroughput sequencing data. Bioinformatics 31:166–169.

48. Mortazavi A Williams BA McCue K Schaeffer L Wold B. 2008. Mapping and quantifying mammalian transcriptomes by RNA-Seq. Nature Methods 5:621–628.

49. Gao B Gupta RS. 2012. Phylogenetic Framework and Molecular Signatures for the Main Clades of the Phylum Actinobacteria. Microbiology and Molecular Biology Reviews 76:66–112.

50. Parks DH Imelfort M Skennerton CT Hugenholtz P Tyson GW. 2015. CheckM: assessing the quality of microbial genomes recovered from isolates single cells and metagenomes. Genome Research 25:1043–1055.

51. Ma H Zeng A-P. 2003. Reconstruction of metabolic networks from genome data and analysis of their global structure for various organisms. Bioinformatics 19:270–277.

